# Computational Design of Myristoylated Cell Penetrating Peptides Targeting Oncogenic K-Ras.G12D at the Effector Binding Membrane Interface

**DOI:** 10.1101/565945

**Authors:** Zhenlu Li, Matthias Buck

## Abstract

A number of small inhibitors have been developed in the recent years to target the cancer driving protein, K-Ras. In this study we propose and design a novel way of targeting oncogenic K-Ras4B.G12D with myristoylated cell penetrating peptides which become membrane anchored and lock the protein into an inactive state. In all atom molecular dynamics simulations such peptides associate with K-Ras4B exclusively at the effector binding region, which, in turn, expected to hinder the binding of down-stream effector proteins (e.g. C-Raf). The myristoylated R9 (Arg_9_) peptide strongly locks K-Ras4B.G12D into orientations that are unfavorable for effector binding. After breaking the cyclic structure and myristoylation, a cell penetrating peptide cyclorasin 9A5, which was designed for targeting the Ras: Raf interface, is also found to be effective in targeting the Ras: membrane interface. The myristoylated peptides likely have high cell permeability due to their mixed cationic/hydrophobic character at the N-terminus, while simultaneously the subsequent multiple charges help to maintain a strong association of the peptide with the K-Ras4B.G12D effector binding lobe. Targeting protein-membrane interfaces is starting to attract attention very recently, thanks to our understanding of the signaling mechanism of an increased number of peripheral membrane proteins. The strategy used in this study has potential applications in the design of drugs against K-Ras4B driven cancers. It also provides insights into the general principles of targeting protein-membrane interfaces.

## Introduction

Our ability to target cancer driving Ras mutations has advanced considerably since 2010, with multiple inhibitors being developed. Fesik and colleagues pioneered methods to search for small molecules that bind to K-Ras in 2012.^1^ In 2013, the first effective inhibitors for K-Ras^G12C^ variant was reported by Shokat and colleagues. The inhibitors rely on the mutant cysteine for binding and act as competitive inhibitors that block GTP binding.^2^ In the same year, an inhibitor targeting the K-Ras: PDE delta interactions was reported by Waldmann and colleagues^3^ and another small inhibitor (Kobe0065) that blocks the Ras: Raf interface was reported by Kataoka and colleagues.^4^ In 2015, Pei and colleagues synthesized a cyclic peptide (cyclorasin 9A5) that targets the Ras: Raf interface.^5^ One year later, Wellspring Biosciences invented a compound ARS-853 (now updated to ARS-1620), which covalently binds to the mutant cysteine of K-Ras^G12C^ in the GDP-bound form.^6,7^ The compound suppresses K-Ras^G12C^ signaling and tumor cell growth.^8^ Overall, well used strategies for inhibiting K-Ras include selectively targeting the cysteine group of K-Ras^G12C^, blocking the membrane localization of Ras or blocking Ras: effector protein interfaces. For the latter, also a monobody, NS1^9^, and a pan-inhibitor, compound 3144,^10^ were recently found effective in blocking the activation of downstream pathways: the extracellular signal-regulated kinase (ERK) pathway or the mitogen-activated protein kinase (MAPK) pathway. While the G12C oncogenic variant is less frequent, the G12D/G12V mutations are commonly found in human tumors, accounting ∼20%-30% of all human cancers.^11,12^ In particular, K-Ras mutations exist in 60% of pancreatic cancer, 17% of lung cancer and 35∼40% of colon cancer.^12,13^ The recent development of promising inhibitors suggests that clinically effective inhibitors towards the Ras protein will be available in the nearer future. Understanding the mechanism of Ras targeting drugs also increases our insight into the critical features of the Ras signaling process itself.

K-Ras signal transduction requires its localization to the plasma- or other cellular membranes, the activation of the GTPase by a nucleotide exchange factor, and once it is in the GTP bound state, binding to effector proteins, such as Raf or PI3K. These events typically happen in a geometrically restricted space at the cytoplasmic membrane. Accordingly, general features of the cell membrane, as well as specific lipid molecules, have significant effects on association, clustering, and on Ras function.^14-16^ Importantly, it is now recognized that in addition to the lipid anchor at the C-terminal hyper-variable region of K-Ras (residues 167 to 185), the folded region of the GTPase (residues 1 to 166) also interacts frequently with the inner leaflet of the plasma membrane. This folded K-Ras region (also called catalytic- or G-domain) can be divided into two functionally different lobes, an effector lobe (res. 1 to 86) and an allosteric lobe (res. 87 to 166).^12^ Both lobes bind to the membrane, based on the results of computational modeling, NMR and FRET.^17-22^ For functional activity via the ERK or the MAPK pathway, the effector lobe needs to associate with down-stream effector proteins, such as Raf and PI3K.^23,24^ Therefore, membrane binding of the effector lobe will occlude the association of an effector protein with this region of the GTPase, and thus attenuate the activity of K-Ras. This finding indicates a possible targeting strategy on K-Ras, by designing inhibitors that lock the proteins into inactive orientations. Recently, Hardy and colleagues found that a compound (Cmpd2) inhibits K-Ras^G12V^ in a lipid dependent manner.^25^ Their follow-up NMR study showed that this compound likely inhibits the function of K-Ras from the membrane surface.^26^

Design of drugs that target the protein-membrane interactions is a relatively new idea. It this study, we propose a strategy of trapping K-Ras onto the membrane using membrane anchored cell penetrating peptides. Cell penetrating peptides are highly cationic small peptides which easily cross the plasma membrane, indicating a great potential for drug delivery into cells.^27-29^ In our prior study of K-Ras, we have found that “off-plane” negative potentials generated by PIP2 at the membrane surface, alter the orientational preference of K-Ras4B, leading to an increased exposure of its effector lobe to the solvent.^20^ As an inference, the opposite, i.e. “off-plane” positive potentials at the membrane surface may effectively also trap the negatively charged effector lobe onto the membrane. Since cell penetrating peptides are highly cationic, it is likely to associate favorably with the effector lobe of K-Ras.G12D. In order to generate an “off-plane” positive potential, the cell penetrating peptides need to be anchored to the membrane, which is accomplished in a straightforward manner by the N-terminal addition of the hydrophobic myristol group. Indeed, a pioneering study by Lee and Tung showed that myristoylated cell penetrating peptides (myristoylated polyarginines n=7-12) have an even higher membrane penetration ability than wild cell penetrating poly-arginines.^30,31^ Excitingly, while the developments of cell penetrating peptides were primarily aimed at enhancing drug transport into the membrane, as noticed above, Pei and collaborators developed a cyclic cell penetrating peptide and found that it directly acts as an effective inhibitor for K-Ras.^5^ The peptide, cyclorasin 9A5, was found to penetrate the membrane easily and efficiently block the Ras: Raf association. The anchoring of myristoylated-cell penetrating peptides as well as their potential to target the Ras: Raf interface, motivated us to test our hypothesis that an “off-plane” positive potential produced by a membrane anchored peptide may lock K-Ras4B into an inactive membrane-bound state. Using an all-atom molecular dynamics study, we find that the myristoylated R9 and a myristoylated linear peptide analogue of cyclorasin 9A5, bind to the effector lobe of K-Ras4B.G12D and thus are expected to attenuate the GTPase’s ability to associate with down-streaming effector proteins. By further optimizing the peptide sequence, the strategy of targeting protein-membrane interfaces may become a potential avenue for future drug design.

## Results and Discussion

### K-Ras4B Binds to the Membrane Doped with Myristoylated Nona-arginine (R9)

Polyarginine (n=7-12), an important type of cell penetrating peptide, has been used to transport biological cargos into a variety of cell.^32,33^ Here, we test nona-arginine (R9) consisting of nine arginine residues in its ability to bind K-Ras by use of computational modeling. The peptide is conjugated with a myristoylate group at its N-terminus (myr_R9) and is inserted into the membrane via this group (Fig. 1a). The simulations were performed firstly at a POPC membrane. Previous simulations have shown that the K-Ras core domain associates with a charge-neutral membrane only transiently. In the presence of myr_R9, the K-Ras significantly binds to the membrane. In the four simulations, the membrane association of the core domain with the membrane was typically established within 100-200 ns (Fig. 1c). Once bound to the membrane, the K-Ras4B maintains the initial membrane bound state during the remainder of the simulations, except in one simulation, where the K-Ras4B core domain dissociates from the membrane at the end of the simulation. In addition, we carried out two additional simulations by initially placing four non-lipidated R9 at the membrane surface. In both simulations, the R9 does not adhere to the membrane and dissociate from the membrane. Occasionally, an escaped R9 binds to the K-Ras4B, however it is not able to bring the core domain of K-Ras4B to the membrane (Fig. S1). Thus, membrane anchoring of R9 is essential for trapping K-Ras4B onto the membrane.

**Figure 1:**
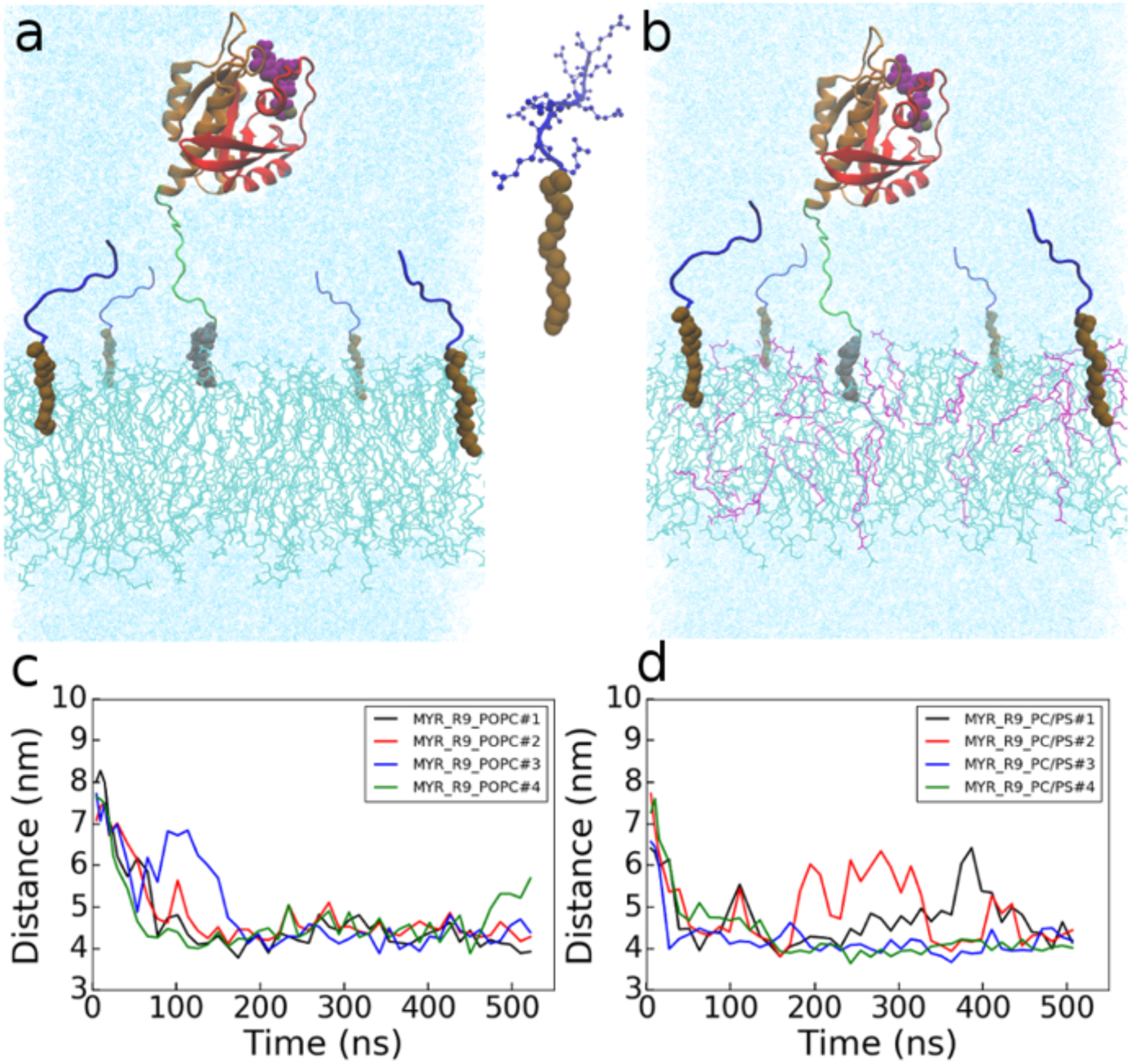
Simulation setup and binding of K-Ras4B to the membrane. Initial structures of K-Ras4B.G12D.GTP: myr_R9 at a (a) POPC and (b) POPC/POPS membrane. The K-Ras4B protein consists of an effector lobe (red, res. 1-86), an allosteric lobe (brown, res. 87 to 166), and the largely unstructured hypervariable region, HVR (green, res. 167 to 185). Proteins shown as mainchain cartoon. R9 peptides are in blue. Small molecules/ions as space filling: farnesyl group (grey); myristoyl group (ochre); GTP (purple); Mg (tan). Membrane shown as lines: POPC (cyan) and POPS (purple). Water is in light cyan. (c, d) plots of the distance between the center of mass of the K-Ras4B core domain and the membrane center as a function of simulation time.

In eukaryotic cells the inner leaflet of the membrane is typically enriched in anionic lipid, such as phosphatidyl serine, PS. K-Ras4A and −4B have been found to bind to such a membrane relatively strongly but still with multiple distinct orientations.^17-22^ Addition of such anionic lipid molecules may neutralize the charge of some of the arginines and thus may increase the binding of myr_R9 peptide residues to the membrane, while possibly decreasing K-Ras – POPS interactions. However, in the simulations where 20% of POPC was replaced with POPS, the binding of the K-Ras4B core domain toward the membrane is also very strong (Fig. 1b and Fig. 1d). In the four simulations, the membrane association of the core domain was established by ∼150 ns (Fig. 1d). In two cases, the K-Ras4B maintains the association once bound to the membrane. In another two simulations, K-Ras4B undergoes dissociation and an orientational adjustment relative to the membrane, but eventually rebinds to the membrane at the end of the simulation. The membrane anchored peptides do not often lie on the membrane surface; thus, the polyArg has a lesser neutralization effect on the negative charge of POPS than anticipated. Therefore, either in a POPC or a POPC/POPS membrane, the core domain of K-Ras4B strongly binds to membrane for 96.9% and 77.7% of the simulation time, respectively, after the first 100 ns.

### Inhibited Orientations of K-Ras4B at the Membrane

In the presence of myr_R9 at the membrane, in all the simulations, the proteins bind to the membrane almost exclusively using the effector lobe (residues 1 to 86) as most of the residues that interact with this cell penetrating peptide are also found in this lobe (Fig. 2a-b). Especially, when anionic POPS lipid molecules are at the membrane, rarely residues of the allosteric lobe are involved in myr_R9 binding (Fig. 2b). Previous computational and experimental studies showed that both the effector and the allosteric lobe interact with the lipid membrane containing 20% of anionic lipid molecules, POPS.^17-20^ In the presence of myr_R9, the effector lobe, which contains the two nucleotide sensitive switch regions, is the dominant region that interacts with the membrane. The binding of K-Ras4B catalytic domain with the membrane is more focused on the effector lobe with the addition of anionic lipid molecules. Fig. 3 shows a typical membrane-binding state of K-Ras4B with the effector lobe in the vicinity of the membrane. Due to the approach of the K-Ras effector lobe to the membrane, the K-Ras4B is not able to access to the C-Raf RBD (RBD-Ras Binding Domain; In Fig. 3a, C-Raf RBD has significant clash with the model membrane). Therefore, myr_R9 effectively locks K-Ras4B into an inactive orientation state.

**Figure 2:**
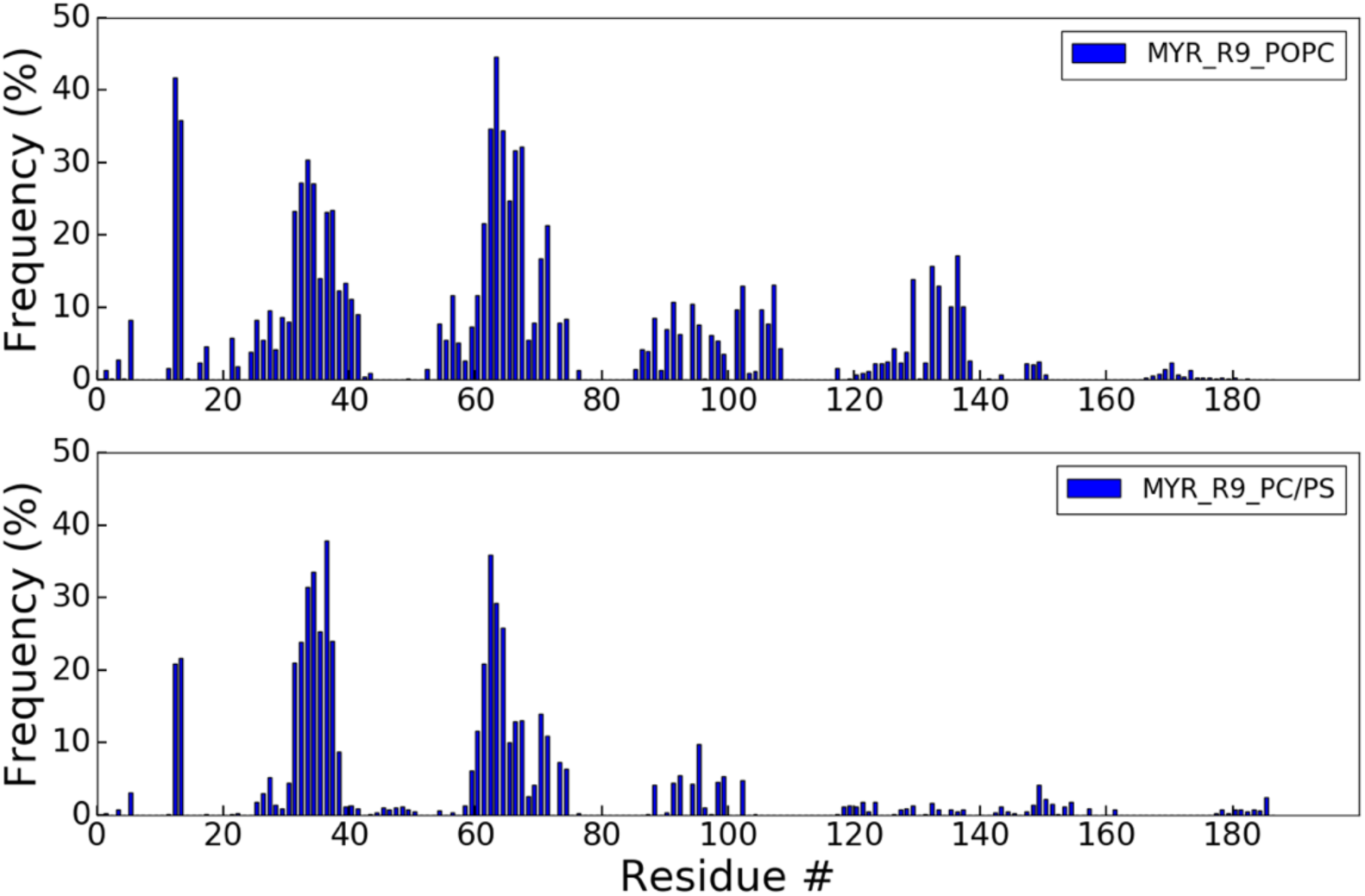
Interface residues of K-Ras interacting with myr_R9. (a-b) Frequency of K-Ras4B: myr_R9 contacts (protein residue atoms within 4 Å of myr_R9 atoms) over the last 400 ns.

**Figure 3.**
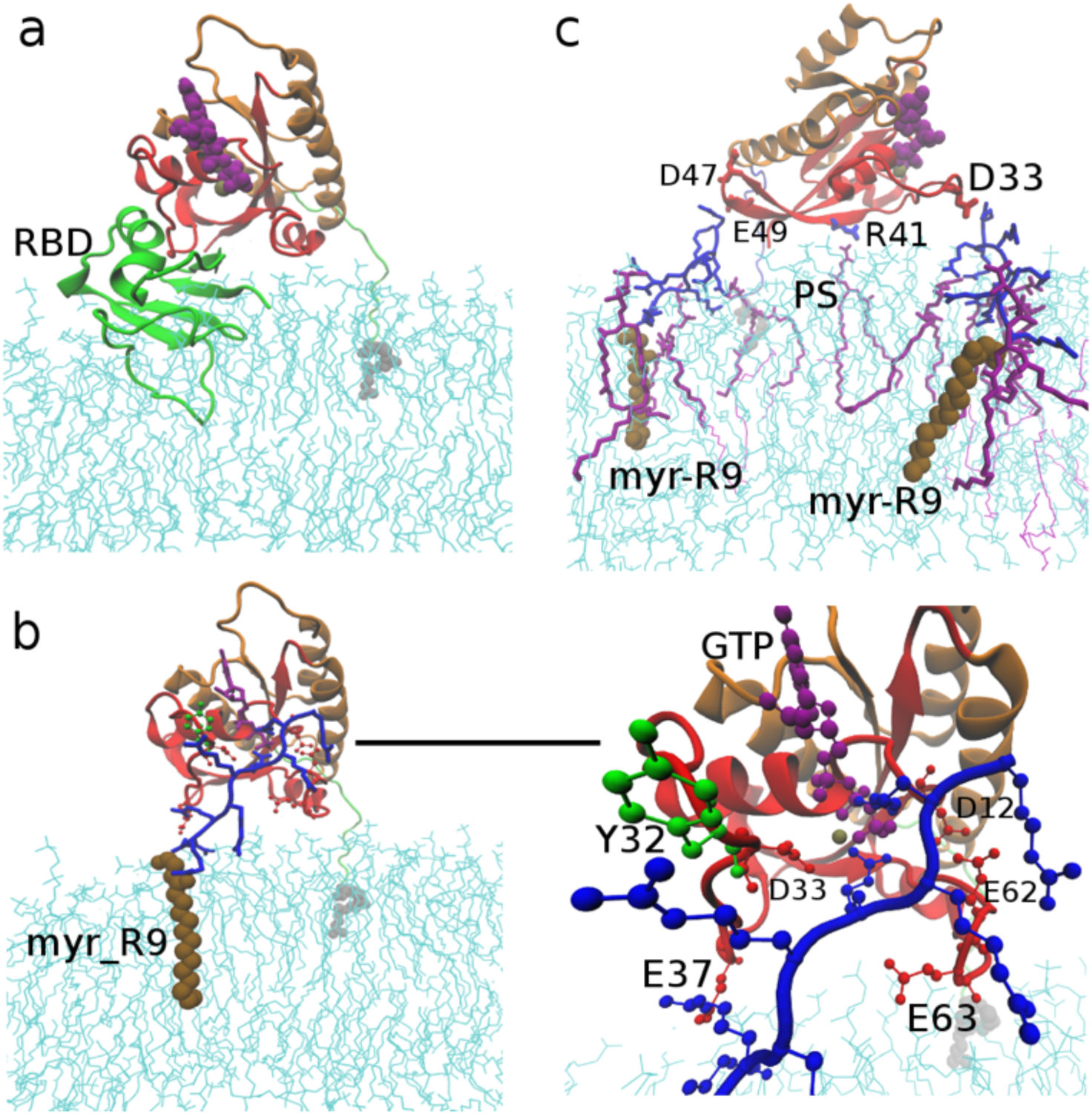
Orientations of K-Ras4B at the membrane that occlude the C-Raf RBD binding site on K-Ras (a) C-Raf RBD: Ras from the X-ray structure [pdb 4G0N, ref. 45] placed at membrane by superposition on the predominant orientation of K-Ras4B sampled in the simulations: as is evident C-Raf RBD has significant clash with the model membrane. Due to the approach of the C-Raf effector domain to the membrane, K-Ras is not able to bind to RBD, being inactive. (b) Left, binding of a myr_R9 towards the K-Ras effector domain at POPC membrane (Snapshot at 500 ns of simulation #3). Right, zoomed in and highlighting the crucial interactions between myr_R9 and K-Ras4B. Electrostatic pairs between Arg and GLU/ASP are dominant. Cation-π interaction between Arg and Tyr32 is also seen and the GTP nucleotide interacts with two arginines of the myr_R9. (c) Interactions of K-Ras4B and two myr_R9 peptides at a POPS membrane (Snapshot at 500 ns of simulation #2). Myr_R9 is surrounded by 3-4 POPS. Color scheme same as Fig. 1.

Looking at the interactions in detail, the K-Ras4B effector domain directly interacts with multiple positive charges of the myr_R9, especially with residues D12, Y32, D33, E37, E62, E63 (Fig. 2a-b). Fig. 3b-c highlights these interactions between myr_R9 and K-Ras4B at a POPC and a POPC/POPS membrane respectively. In the first example (Fig. 3b), five electrostatic pairs are established between the myr_R9 and the K-Ras4B. It is noticeable that two arginines from myr_R9 are bound to the phosphate group of GTP. In the second example (Fig. 3c), K-Ras4B is bound to two myr_R9 peptides. In addition, K-Ras residue R41 associates with a POPS lipid molecule. Many of the arginines of myr_R9 are surrounded by 3 or 4 POPS lipid molecules and are thus partially neutralized by these lipids. However, the C-terminal arginine(s) could easily access to the negative charged residues of K-Ras catalytic region. Importantly, the mutated residue, D12, which makes K-Ras4B oncogenic, was not involved in membrane binding in prior simulations.^20^ This is because, D12 is spatially distant from the membrane anchored HVR region and is hard to get to the membrane surface when the preferred orientation states are sampled.^20^ Since myr_R9 does not lie on the membrane surface horizontally, the arginine sidechains could access even distant regions of K-Ras, such as the location of D12. The attraction between D12 and arginine also brings the region near D12 closer to the membrane. Given that the K-Ras G12D mutation is a major oncogenic form of K-Ras, binding between D12 and the myr_R9 may be selective in inhibiting the activity of this oncogenic mutant, but further calculations and experiments will be necessary to confirm this.

Fig. 4 compares the orientation distributions of K-Ras relative to the membrane in the current simulations to the distributions in prior simulations of the highly homologous K-Ras4A at a POPC/POPS membrane and at a POPC/PIP2 membrane.^20^ At a POPC/POPS membrane, K-Ras could evenly sample orientations of the effector lobe (O1; *β*1-*β*3 and *α*2; residues 1 to 74) or the allosteric lobe (O3/O4; *α*3-*α*5; residues 87-166), or occasionally parts of both lobes (O2; *α*2-*α*3; residues 66-104). In state O1 and O2, the effector lobe is not available for effector protein binding. However, at a POPC/PIP2 membrane, K-Ras samples only one dominant orientation, with the allosteric lobe binding to the membrane and effector lobe is exposed to solvent (O3) and is thus predicted to increase K-Ras activity by easily allowing effector protein binding.^20^ In the presence of myr_R9, at a POPC or POPC/POPS membrane, the orientation distribution is mostly close to the O1 state, and less so to the O2 state. In all these orientations, the K-Ras effector lobe is unavailable for binding of the C-Raf RBD, which is known to form the tightest interaction between these two protein regions. The orientations are not exactly the same as seen with the simulations at a POPC/POPS membrane. This is because, in the presence of myr_R9, distant residues such as D12 also get closer to the membrane, and this typically lifts up the membrane binding region on the other side of the protein, i.e. the allosteric lobe of K-Ras4B.

**Figure 4:**
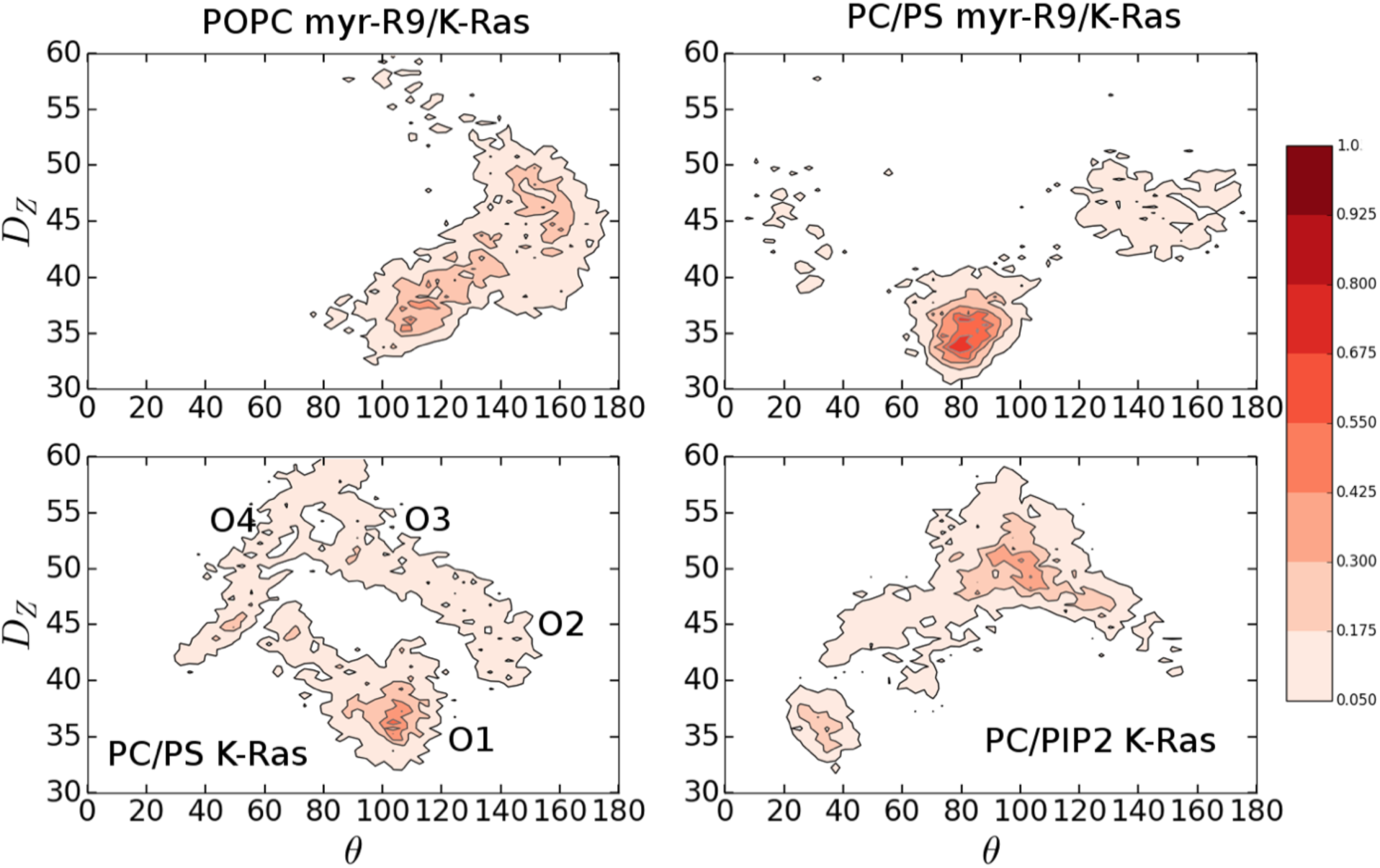
Orientations of K-Ras4B at the membrane. Contour maps of orientation parameters D_z_ and θ, sum-averaged over the four simulations. The bottom panels are results from our prior study of K-Ras at POPC/POPS and POPC/PIP2 membrane, in the absence of myr_R9.^20^ See methods for definition of the variables (D_z_ and θ). The probability is calculated by dividing the number of frames a structure exists in an orientation state by the total number of sampling frames. The probability is scaled to 1 for the maximally populated state in simulations of myr_R9 at ta POPC/POPS membrane.

### Electrostatic Interaction as a Major Factor for Effector Lobe: myr_R9 Association

The binding of the effector lobe to the membrane is largely affected by electrostatic interactions. The effector lobe has 15 negatively charged residues and 6 positively charged residues, while the allosteric lobe has 13 negative residues and 15 positive residues. On average, then, the effector lobe has a negative electrostatic potential, by contrast to the allosteric lobe. This is the predominant reason why the effector lobe binds strongly to the Ras Binding Domain (RBD) of the effector protein C-Raf, as this domain possesses a highly positively charged electrostatic potential surface. Electrostatic residue interaction pairs between the effector lobe of K-Ras and the C-Raf RBD includes E37:R66; E31/D33:K84 and D38:R89. In the presence of myr_R9, Ras residues E33 and E37 are strongly bound to the membrane anchored peptide, thus these interactions must disrupt the association between C-Raf RBD and K-Ras. As indicated in our previous study, PIP2 lipids generate a broad and partially “off-plane” negative potential at the membrane, therefore, compared to a POPC/POPS membrane, a POPC/PIP2 membrane will push the effector lobe of K-Ras4B away from the membrane-solvent interface.^20^ By contrast, the myr_R9 peptide overall points away from the membrane surface (Fig. 5a). The C-terminus of R9 is about 2.5 nm to 3.0 nm away from the membrane center (0.5-1.0 nm away the membrane surface). The addition of POPS only slightly enhances the binding of the myr_R9 to the membrane surface. Due to the deviation of myr_R9 from the membrane surface, the charge distribution of the myr_R9: membrane system broadens (Fig. 5b). The distribution of positive charges coming away from the membrane surface generates an “off-plane” positive potential. Therefore, as expected, the effector lobe with multiple negatively charged residues is attracted by these positive charges, and is consequently strongly bound to the membrane. In addition, the myr_R9 peptide does not interact with the polybasic HVR of K-Ras4B. The distance between the HVR and peptide is mostly larger than 2 nm (Fig. 5c). This is because the HVR has multiple positive residues and thus repels the arginines in myr_R9. In this sense, the peptides may have additional mechanisms by which they inhibit K-Ras4B, as Ras signaling is known to involv higher order oligomers or clusters of K-Ras and effector proteins.^14-16^ The repulsion between myr_R9 and the HVR may keep K-Ras4B proteins apart, if these peptides localize between K-Ras proteins. This will dramatically attenuate the number of clusters and thus the signaling of K-Ras.

**Figure 5:**
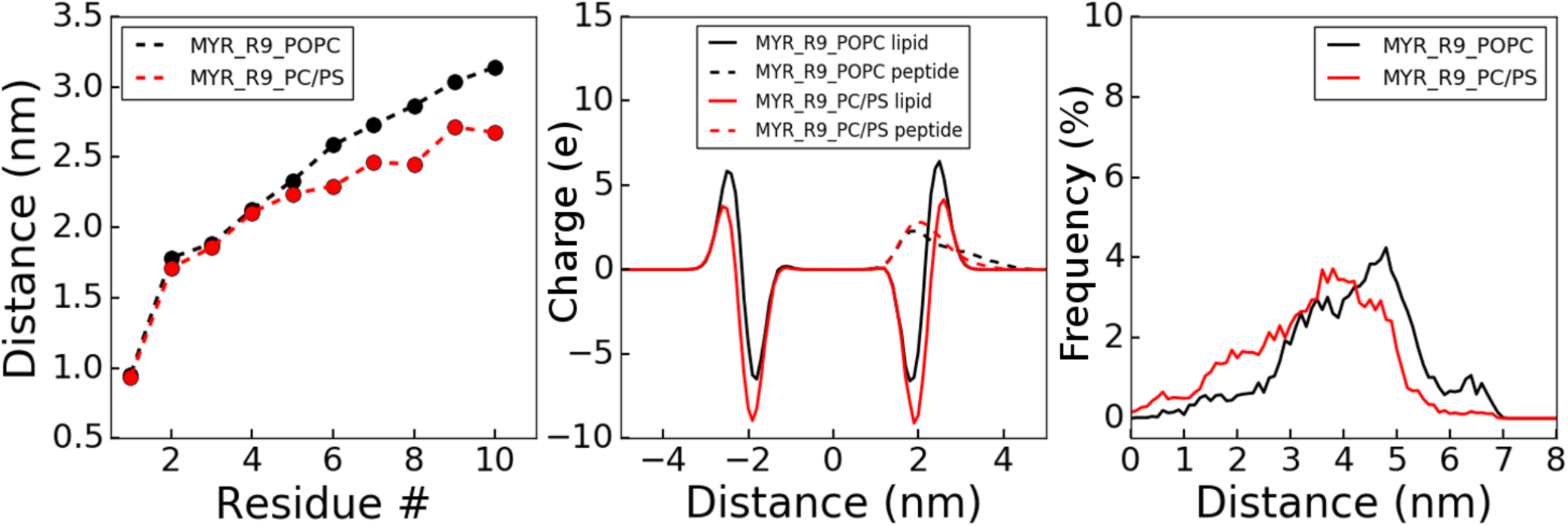
Distances and Electrostatics involved in K-Ras4B: myr_R9 binding. Averaged over four simulations. (a) Distance of individual residues to the center of mass of the membrane; (b) Distribution of charge across the model membrane (contribution from lipid molecules as well as peptides); (c) Distribution of distance between the center of mass of myr_R9 and the center of mass of the K-Ras4B HVR in the two membrane systems.

We further carried out two additional tests: a) by substituting the myristoyl group with a farnesyl group and separately, b) by replacing the R9 with an amino acid sequence (RKTFLKLA) derived from residues 143-150 of the C-Raf CRD, as this segment was also found to interact with K-Ras in solution.^34^ In the computational study, we made a replacement of a myristoylate with a farnesyl (See Methods). In the presence of far-R9, K-RAS4B was found to have even stronger membrane binding. Remarkably, K-Ras4B binds to membrane with the effector lobe exclusively (Fig. S2). The reason for this is unclear. By replacing the R9 with the CRD segment (quite unlike the cell penetrating peptide in character), the simulations showed the peptide has no locking effect of the K-Ras4B orientation (Fig. S3). This is likely because it has relatively less charge compared to the two other peptides tested in this study. Furthermore, the peptide is partially inserted into the membrane with its phenylalanine and leucine sidechains. Therefore, it cannot access distal regions of K-Ras4B. These two trial simulations indicate that the anionic character of cell penetrating peptides as well as the presence of a membrane anchor are both crucial for their capability to associate with the effector lobe.

### A Linear Myristoylated Cyclorasin 9A5 Variant Effectively Traps K-Ras4B into an Inactive State

Cyclorasin 9A5 is a cyclic cell penetrating peptide with a sequence of -Ala-Thr-Trp-Gln-Nle-Phe-Arg-Nal-Arg-Arg-Arg-.^5^ The peptide can permeate the plasma membrane and block interactions at the Raf binding interface of Ras. In order to anchor the peptide to the membrane, the cyclic peptide was broken before the N-terminal Ala and that Ala was mutated to Gly and N-terminally myristoylated (Fig. 6a). Then myristoylated linear peptides were added to a POPC or POPC/POPS membrane. Overall, the peptide has less positive charges than R9. Accordingly, it has less binding toward the K-Ras4B effector lobe. Bound to a POPC membrane, myr_cyclorasin 9A5 has rare interactions with K-Ras4B (Fig. S4). However, when anionic lipid molecules are added to the membrane, the K-Ras4B core region became attached to the membrane (Fig. 6b), although the overall percentage of time when it is membrane-bound is reduced, compared to the myr_R9 containing membrane (Fig. 6c, in comparison to Fig. 2). After 100ns, the membrane bound state of the Ras catalytic domain exists for 55% of the remaining simulation time.

**Figure 6:**
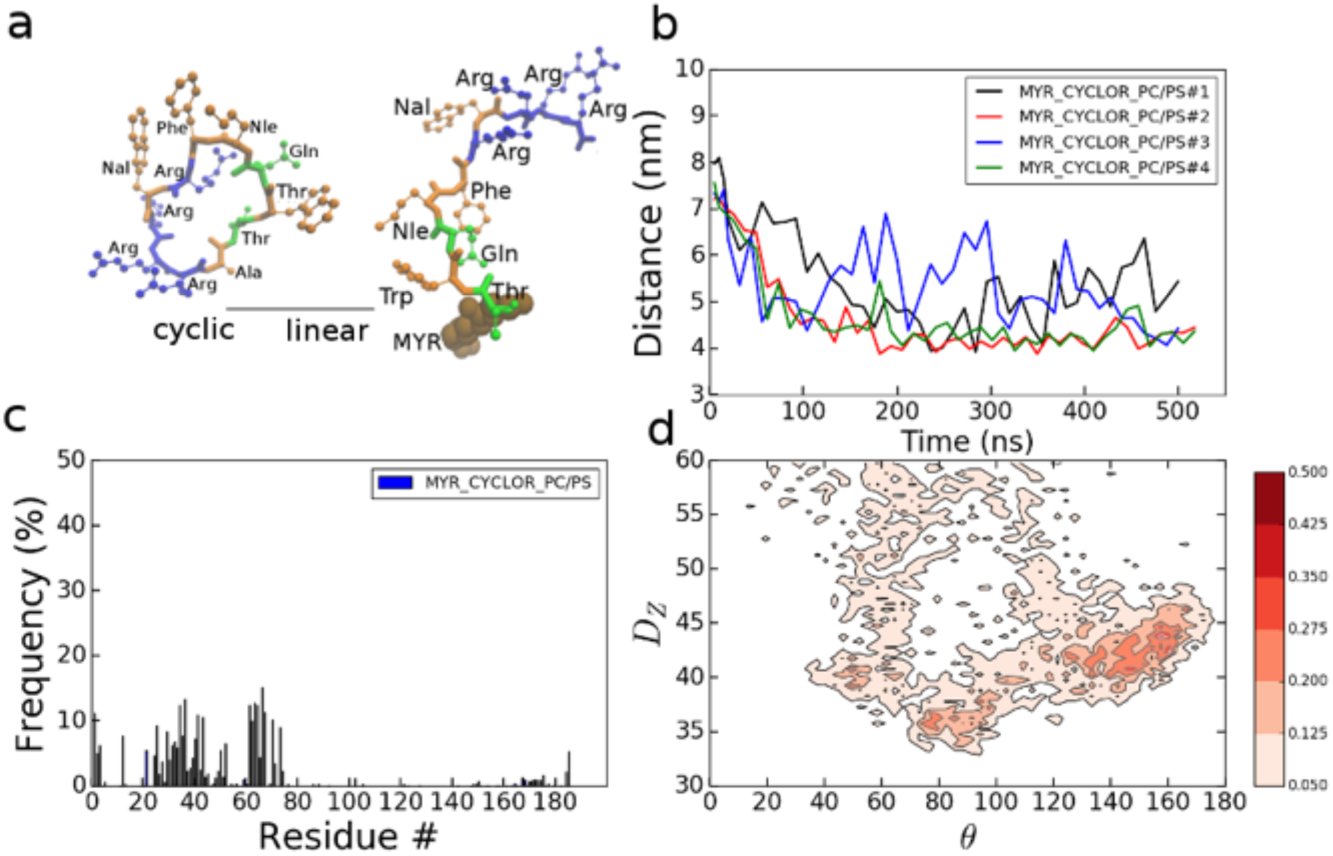
Association of K-Ras4B with myr_cyclorasin 9A5 at a POPC/POPS membrane. Initial structure of myr_cyclorasin 9A5, left cyclic and right, linear + myr. (b) Time evolution of distance between the center of mass of the K-Ras4B core domain and the membrane center. (c) Contact frequency of K-Ras4B residues with myr_cyclorasin 9A5. (residues within 4 Å of myr_cyclorasin 9A5). (d) Contour maps of orientation parameters D_z_ and θ that are popular as sum of the four simulations and scaled as above.

Nevertheless, it appears that myr_cyclorasin 9A5 has a “locking” effect similar to that of myr_R9 on the orientations of K-Ras4B at a POPC/POPS membrane. Analysis of the K-Ras4B: myr_cyclorasin 9A5 interface showed that the arginine residues (7, 9, 10, 11) of myr_cyclorasin 9A5 interact with the residues D12, R41, E62, E63, Y64 of K-Ras4B (Fig. 6b). Furthermore, based on the results of Pei and colleagues, the cyclic cyclorasin 9A5 mainly binds to the switch I region of Ras including residues I24, Q25, D33, E37, S39, and a small pocket between switch I and switch II including residues L56, D57, M67, R73, T74, G75, L79.^5^ In the linear form, some of the membrane headgroups and peptide residues are also close to these residues, while many of the interactions are not the same. Despite this, the linear peptide makes many contacts, suggesting that compared to the cyclic peptide, it inhibits the activity of K-Ras4B via a different mechanism by locking it into an inactive orientation relative to the membrane, rather than by blocking effector protein binding directly. The orientation distribution of K-Ras4B is similar to the results of simulations of K-Ras4B at a POPC/POPS membrane in the presence of the myr_R9 peptide (Fig. 6d), however the strength of K-Ras: membrane association appears to be reduced. In order to achieve a stronger membrane binding, probably additional arginines need to be added to the peptide. In comparison to myr_R9 (Fig. 5a), the C-terminus of cyclorasin 9A5 is much closer to the membrane (Fig. 7a). The peptide roughly lies on the membrane surface interacting with membrane headgroups. A membrane adhesion of the cyclorasin is largely assisted by aromatic residues Trp and Phe (Fig. 7a). Similar to the myr-R9, on one hand, the linear cyclorasin 9A5 generates an additional positive charge distribution and thus helps to attract the effector lobe of K-Ras4B (Fig. 7b). On the other hand, myr_cyclorasin 9A5 is distant from the polybasic HVR of K-Ras4B, due to electrostatic repulsion (Fig. 7c).

**Figure 7:**
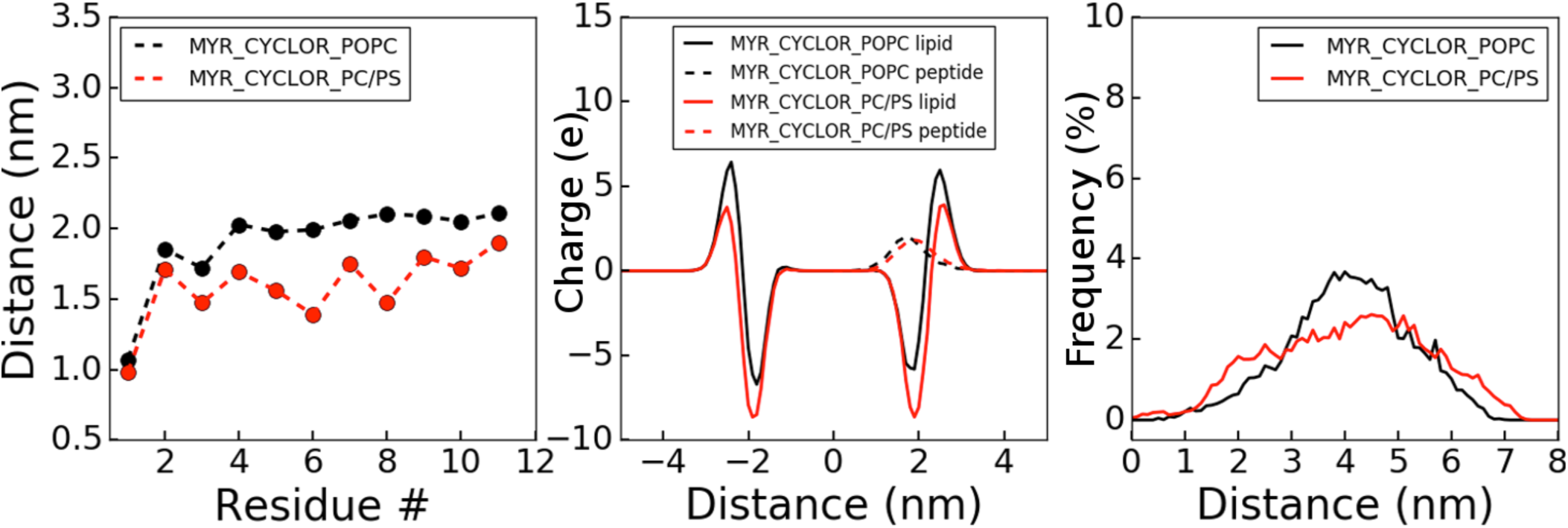
Electrostatics in K-Ras4B: myr_cyclorasin binding. (a) Distance of individual residues of myr_cyclorasin 9A5 to the center of mass of the membrane; (b) distribution of charge across the model membrane (contribution from lipid molecules as well as peptides); (c) distribution of distance between the center of mass of myr_cyclorasin 9A5 to the center of mass of the K-Ras HVR.

## Conclusion

In this study, we propose a scheme for targeting protein-membrane interactions. We use a membrane-anchored peptide to trap an oncogenic membrane-resident protein, K-Ras4B G12D, into an inactive orientation at the membrane with respect to binding of effector proteins. The lipidation/membrane anchoring of these peptides helps to generate an “off-plane” positive potential, which attracts the K-Ras effector lobe onto the membrane surface. There are advantages in targeting the protein-membrane interaction with membrane bound therapeutics. First, effectively confining molecules into two dimensional space increases their local concentration at the membrane. Second, due to the limited short length, such molecules could selectively bind to only certain surfaces of a membrane peripheral protein, thus affecting its membrane bound configuration. Third, with multiple arginines and a lipid anchor, the peptide could be transported to and anchored into the inner leaflet of cell membrane easily. However, the addition of excessive positive charges at the membrane may also affect the other cellular processes or create membrane pores, which could be good for killing cancer cells, but harmful for normal cells.^35^ In future studies, the target specificity and selectivity of the peptides should be examined experimentally. A further optimized design of a lipo-peptide may become an efficient way of targeting K-Ras. Fig. 8 shows possible features that may improve the lipo-peptides in targeting K-Ras4B. Firstly, multiple arginines are needed to maintain both the cell penetration ability and the high affinity toward the K-Ras effector lobe. In order to obtain a better selectivity toward K-Ras, the lipid anchor may be replaced by other types of groups that directly bind to or cluster with the farnesyl group of K-Ras. Different Ras isoforms are laterally segregated into spatially distinct nanodomains.^14^ Different from the saturated lipid anchor-myristoyl group, the unsaturated lipid anchor-farnesyl group, prefers lipid-disordered (l_d_) membrane domain which is abundant in unsaturated lipid molecules.^16,36^ Therefore, lipid anchors such as farnesyl that likes to localize into lipid disordered (l_d_) domains possibly enhance the selectivity towards K-Ras. Besides, a small segment that may specifically recognize individual or multiple residues, for example the G12D, G12C or G12V mutation of K-Ras4B, should largely enhance the binding specificity toward a particular K-Ras mutant. Thus the peptide will have a unique binding towards K-Ras4B and in the meantime effectively locks the K-Ras into an inactive state.

**Figure 8:**
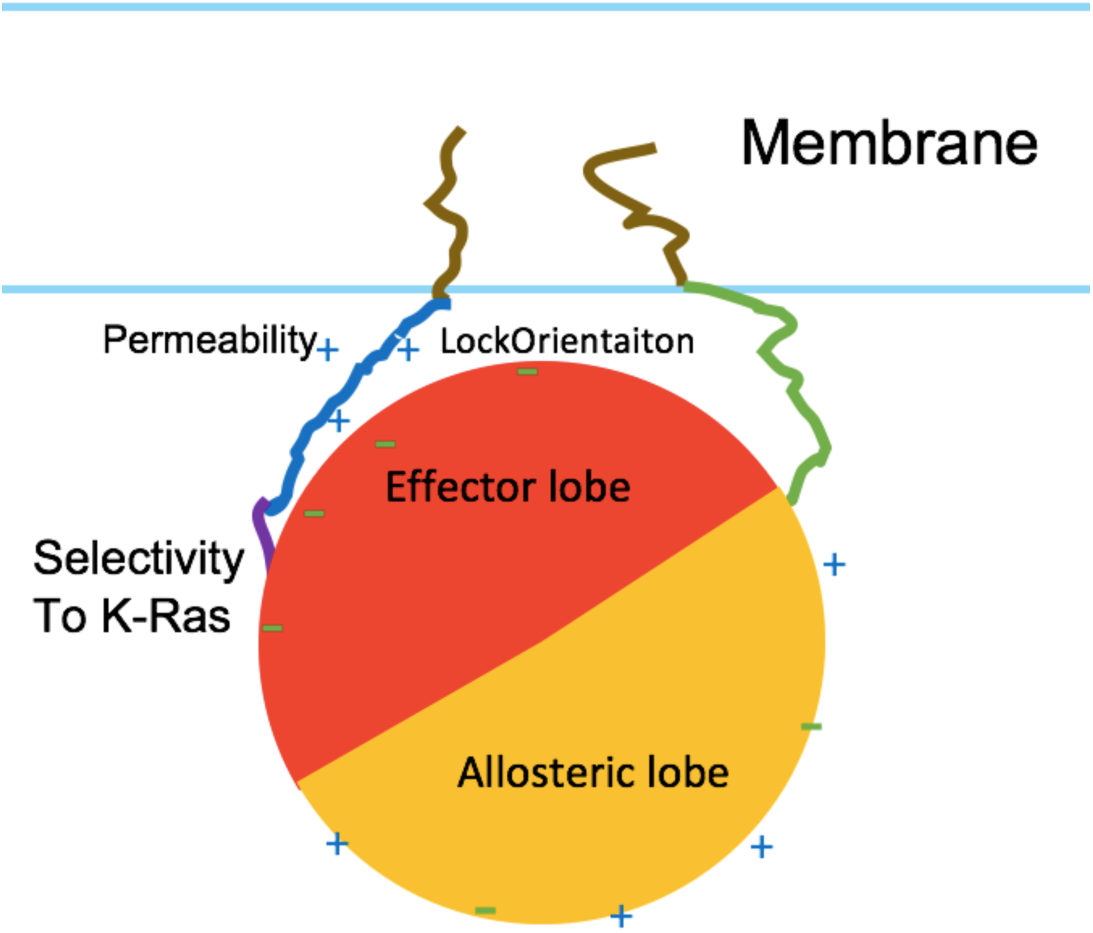
Schematic picture of an ideal design of an inhibitor that targets K-Ras at the membrane. Lipid anchor (ochre) selectively binds to farnesyl group (top to the right) of K-Ras4B; Polybasic peptide (blue) binds to effector lobe and locks K-Ras onto the membrane; a segment (purple) specifically recognizes the K-Ras4B.G12D mutant.

## Methods

### Simulation set up

The membrane consisted of 300 POPC (1-Palmitoyl-2-oleoyl-sn-glycero-3-phosphocholine) or 284 POPC /64 POPS (1-palmitoyl-2-oleoyl-sn-glycero-3-phospho-L-serine) molecules. The number of anionic lipid molecules are equal in each monolayer of the membrane. In order to maintain a balanced lateral pressure of the membrane, two POPC lipid molecules were removed from the leaflet where the peptide and K-Ras4B are anchored. The model membranes were created by the CHARMM-GUI^37^ and equilibrated for 100 ns before adding peptides and K-Ras4B onto the bilayer. Both myr_R9 or -cyclorasin 9A5 were relaxed via a simulation of 20 ns in solvent before placing them at the membrane. After the relaxation, the myr-groups of four myristoylated peptides were partially pre-inserted into the membrane, leaving the rest of the peptide roughly perpendicular to the membrane (full myr group insertion occurred quickly at the start of the simulation). Therefore, the great majority of peptide residues do not have contact with the membrane at the beginning of the simulation. The four peptides are separated from each other by placing them at the corners of membrane/solvent periodic boundary box. The GTPase K-Ras4B was anchored to the membrane but via a farnesyl group. The crystal structure of G12D K-Ras4B (PDB ID: 4DSO) was used for modeling K-Ras4B.^38^ The crystal structure has a Magnesium ion and has a non-hydrolysable GMP-PCP as the bound nucleotide, which was changed to GTP in the simulations. The protein-membrane system was solvated by TIP3P water in a simulation box of about 110×110×170 Å^3^, setting up periodic boundary conditions. Sodium and Chloride ions were added to a near-physiological concentration of 150 mM and to make the system charge neutral. In total, the simulation box consisted of about 155,000-190,000 atoms.

### Simulation parameters

The myristoyl- and farnesyl groups were parameterized via CHARMM-CGEnFF.^39^ In order to add a farnesyl group, a cysteine residue is added to the R9 peptide and the farnesyl is covalently linked to this cysteine residue. The cyclorasin 9A5 has a sequence of -Ala-Thr-Trp-Gln-Nle-Phe-Arg-Nal-Arg-Arg-Arg-, where Nle is norleucine and Nal is D-ß-naphthylalanine. Parameters of residues Nle and Nal in cyclorasin 9A5 were also produced via CHARMM-CGEnff. CHAMRM36m force field was used in the simulations for water and biomolecules.^40^ CHARMM36m has made corrections on the guanidinium interactions between ARG and GLU/ASP, reducing their electrostatic interactions. Even though the potential correction does not affect protein-membrane interactions directly, it has an indirect effect on the orientation preference of K-Ras relative to the membrane.41 The van der Waals (vdW) potential was truncated at 12 Å and smoothly shifted to zero between 10 and 12 Å. The Particle-Mesh Ewald (PME) method was used for calculating the long distance electrostatic interactions. The SHAKE algorithm was applied for all covalent bonds to hydrogen. A time step of 2 fs was employed and neighbor lists updated every 10 steps. The temperature was coupled by to a Langevin thermostat of 310 K, whereas the pressure control was achieved by a semi-isotropic Langevin scheme at 1 bar.

Four independent simulations were performed for myristoylated-R9 at a POPC and a POPC/POPS membrane, or for myristoylated linearized cyclorasin 9A5 at a POPC/POPS membrane. Farnesylated R9, myristoylated C-Raf 143-150, or myristoylated cyclorasin 9A5 was also simulated at a POPC membrane, with two independent runs. Each individual simulation was first started with a harmonic constraint on the protein Carbon alpha atoms (force constant of 1 kcal/mol*Å^2^) for 1 ns. All systems were simulated and equilibrated for 20-30 ns using the NAMD/2.12 package.^42^ The production simulations were performed for 500 ns on the Anton supercomputer, which is optimized for molecular dynamics.^43^

### Analysis

The trajectories were analyzed with VMD^44^ and with scripts for standard analysis such as the distance between two groups. In analyzing the simulations, in most cases, data after the first 100 ns were used unless stated otherwise. The variables D_z_ and θ were defined to describe the orientations of K-Ras4B relative to the membrane in the same way as in our prior study.^20^ Variable D_z_ was determined by the distance (at Z direction) between the center of mass of the effector lobe and the center of mass of the membrane. The different orientations of Ras relative to the membrane originate from the rotation about a main axis. A direction vector (Vy) denoting the average orientation of *β*4 and *β*5 in Ras is chosen as the direction of main axis, and a second direction vector Vx is defined as the vector that connects the center of mass of effector lobe and that of allosteric lobe. The principal axis Vz is then determined by the cross product of vector Vx and Vy. The cross angle between the normal direction to the membrane bilayer plane (Mz) and the direction vector Vz determines variable θ, which describes the tilt of the Ras core domain relative to the membrane.

## Acknowledgements

This work was supported by NIGMS grant R01GM112491 to the Buck laboratory and used the Extreme Science and Engineering Discovery Environment (XSEDE) Stampede at theTexas Advanced Computing Center (TACC), the Ohio Supercomputer Center (OSC), as well as local computing resource in the core facility for Advanced Research Computing at Case Western Reserve University. Anton Computer time was provided by the Pittsburgh Supercomputing Center (PSC) through Grant R01GM116961 from the National Institutes of Health.

