## Supporting_Information for "Computational Design of Myristoylated Cell Penetrating Peptides Targeting Oncogenic K-Ras.G12D at the Effector Binding Membrane Interface"

Zhenlu Li[1*] and Matthias Buck[1,2,3,4*]

[1]Department of Physiology and Biophysics, Case Western Reserve University, School of Medicine, 10900 Euclid Avenue, Cleveland, Ohio 44106, U. S. A.

[2]Department of Pharmacology; [3]Department of Neurosciences; and [4]Case Comprehensive Cancer Center, Case Western Reserve University, School of Medicine, 10900 Euclid Avenue, Cleveland, Ohio 44106, U. S. A.

Figure S1: Snapshots for simulation of K-Ras at a POPC membrane doped with R9 peptides (no myr anchor).

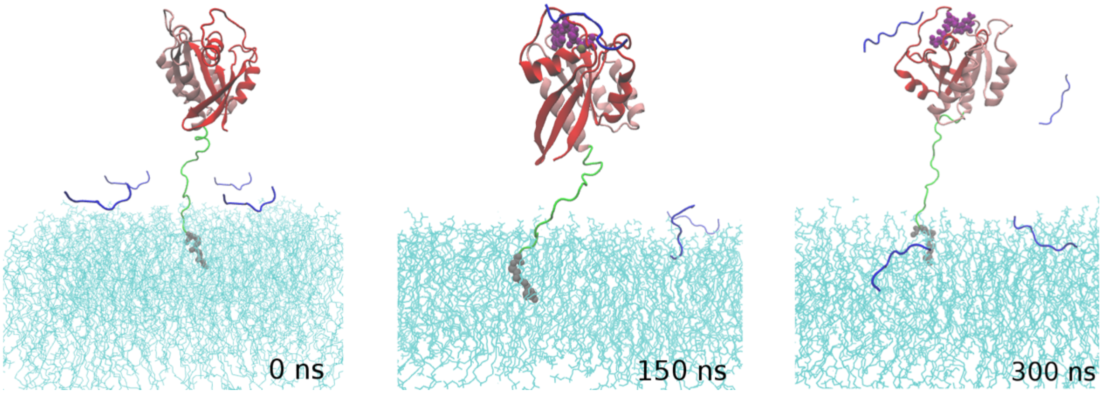


Figure S2: Simulation of K-Ras at a POPC membrane doped with far_R9. (left) Time evolution of distance between the center of mass of the K-Ras4B core domain and the membrane center. (middle) Frequency of K-Ras4B: myr_R9 contacts (protein residue atoms within 4 Å of myr_R9 atoms) over the last 300 ns simulations. (right) Contour maps of orientation parameter $D_z$ and θ.

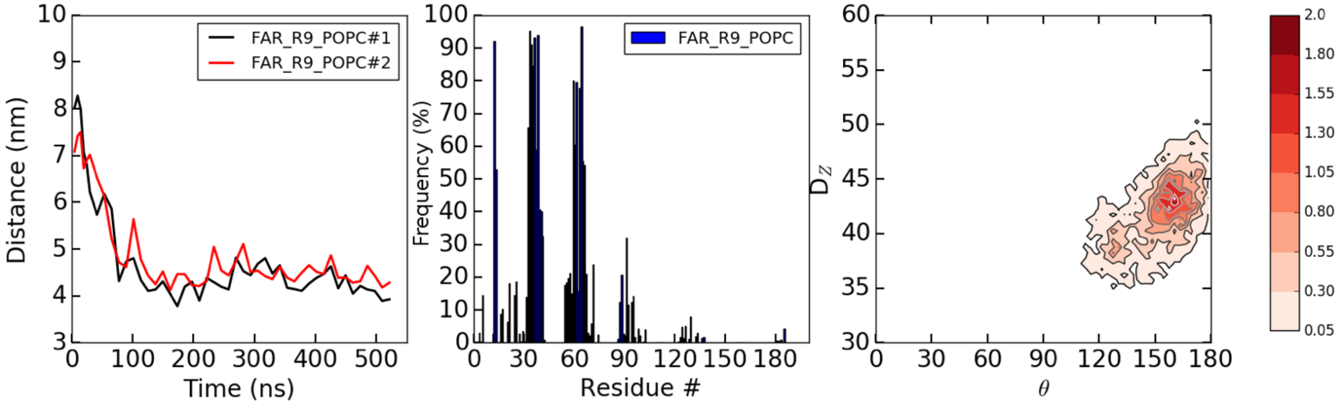

Figure S3: Simulation of K-Ras at a POPC membrane doped with myr_CRD peptide (143-RKTFLKLA-150). (left) Time evolution of distance between the center of mass of the K-Ras4B core domain and the membrane center. (middle) Frequency of K-Ras4B: myr_CRD peptide contacts over the last 300 ns simulations. (right) Contour maps of orientation parameter $D_z$ and θ.

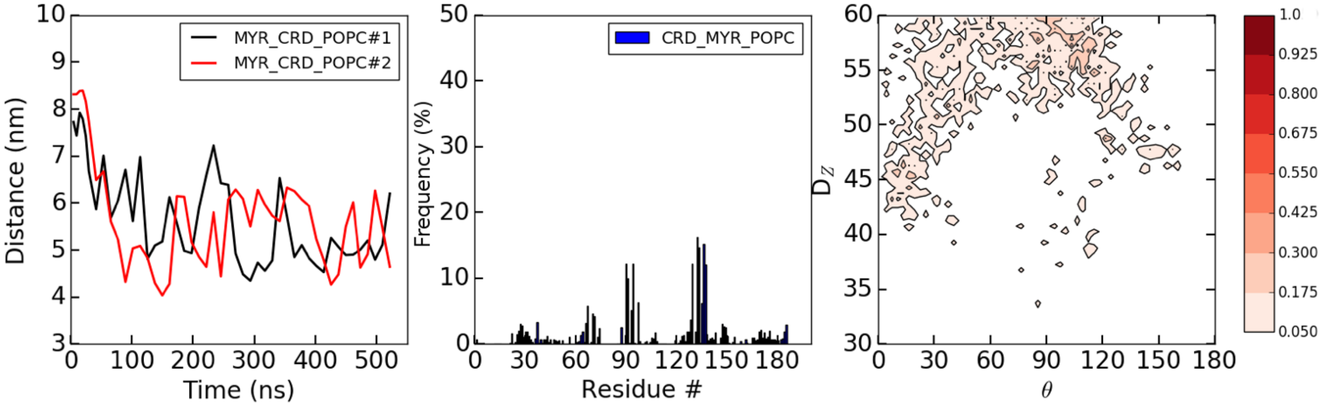


Figure S4: Simulation of K-Ras at a POPC membrane doped with myr_Cyclorasin. (left) Time evolution of distance between the center of mass of the K-Ras4B core domain and the membrane center. (middle) Frequency of K-Ras4B: myr_Cyclorasin contacts over the last 300 ns simulations. (right) Contour maps of orientation parameter $D_z$ and θ.

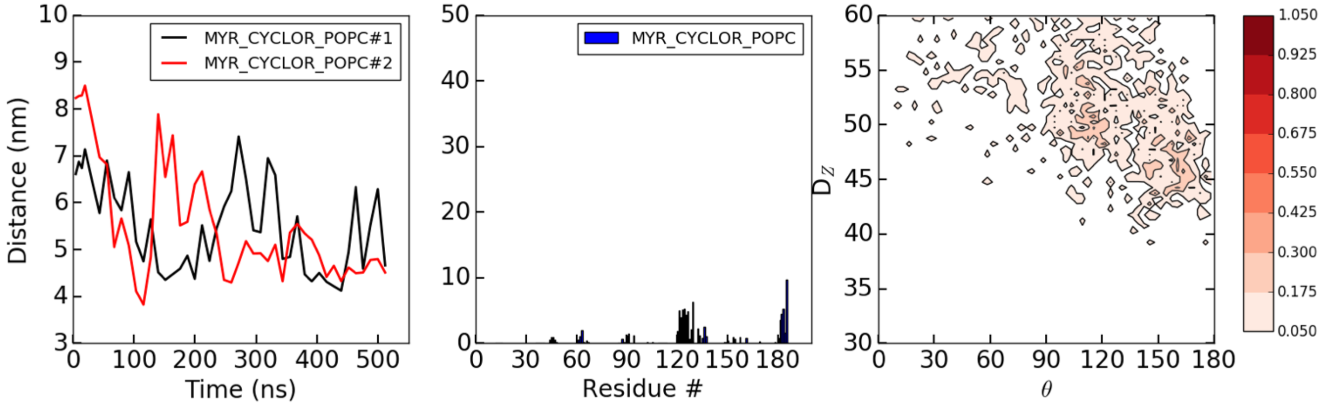
